# Exact switching time distributions for autoregulated gene expression models with mRNA and protein descriptions

**DOI:** 10.1101/2024.06.24.600551

**Authors:** Shan Liu, Bingjie Wu, Chen Jia

**Affiliations:** Applied and Computational Mathematics Division, Beijing Computational Science Research Center, Beijing 100193, China

## Abstract

In this study, we obtain the exact switching time distributions between the free and bound gene states for a detailed stochastic model of an autoregulatory genetic feedback loop with transcription, translation, mRNA and protein decay, as well as protein-gene interactions. The analytical solution generalizes and corrects the previous ones obtained in [Phys. Rev. Lett. 101, 118104 (2008)] and [Nat. Commun. 9, 3305 (2018)] for a reduced model of an autoregulatory loop that ignores the mRNA dynamics. We find that when the mRNA dynamics is modelled explicitly, the holding time in the free gene state can produce three shapes of steady-state distributions (decaying, bell-shaped, and bimodal). In particular, the detailed model with both mRNA and protein descriptions can produce a distribution shape that the reduced model fails to capture — the detailed model of a negative (positive) feedback loop can display a bimodal (bell-shaped) holding time distribution, while the reduced model cannot. Interestingly, we also find that an autoregulatory loop can produce a heavy-tailed holding time distribution and the origin of this heavy-tailed phenomenon is clarified using our analytical solution. Finally, we investigate how the distribution shape is affected by the type of feedback, the binding and unbinding rates, and the transcription rates.

## 1 Introduction

In living cells, the expression of a gene is often controlled by that of other genes, forming various gene regulatory networks [1]. Autoregulation, whereby the protein expressed from a gene binds to its own promoter and activates or suppresses its own transcription, is one of the most common network motifs [2]. Both experiments and theory have revealed how autoregulation modulates fluctuations in the gene product numbers [3–5], the shape of the gene product number distribution [6, 7], as well as the response speed of transcription networks [8, 9].

Over the past two decades, numerous strides have been made in the modelling and theory of autoregulatory gene circuits. A complete stochastic model of autoregulation should include transcription, translation, as well as protein-gene interactions [10]. However, for simplicity and analytical tractability, some studies ignore the mRNA dynamics and consider a simplified model with only the protein description [10–13]. In the literature, various simplified models of autoregulation have been proposed based on different implicit assumptions. Among these models, some assume that protein molecules are produced one at a time [11, 13], while others assume that protein molecules are produced in stochastic bursts [10, 12]. Another distinguishing feature is that some models ignore protein-gene binding fluctuations and assume that there is no change in the protein number when a protein binds to a gene or when it unbinds [11, 12], while other models take such fluctuations into account and assume that the protein number decreases by one when a protein binds to a gene and increases by one when unbinding occurs [10, 13]. In the former case (binding fluctuations are ignored), the chemical master equation (CME) describing an autoregulatory loop can be solved analytically both in steady state [11, 12, 14, 15] and in time [16, 17]. However, in the latter case (binding fluctuations are taken into account), only the steady-state solution has been obtained [10, 13] and the time-dependent solution is only obtained in the special case of fast gene state switching [18].

Due to protein-gene interactions, the gene of interest in an autoregulatory loop can exist in either a free (unbound) or a bound state [13]. While the exact protein number distribution in an autoregulatory circuit has been studied extensively, the holding time distributions in the free and bound gene states have received considerably less attention. Experiments have shown that holding times in different gene states play a significant role in various cellular processes, such as transcriptional regulation [19], cell differentiation [20], epigenetic modifications [21], and response to environmental changes [22, 23]. For example, an approximately deterministic holding time with very little noise may result in periodic switching between gene states and may further drive robust biological oscillations [24]. In previous studies [25, 26], the holding time distributions in the free and bound states have been obtained in closed form in a simplified model of autoregulatory loops that ignores the mRNA dynamics. The difference between Refs. [25] and [26] is that the former ignores protein-gene binding fluctuations, while the latter takes such fluctuations into account. Thus far, the switching time distributions have not been solved exactly in a detailed model of autoregulatory circuits with both mRNA and protein descriptions and the corresponding distribution shapes still remain unknown. Elucidating and understanding such behaviour is important since it helps reveal the nature of biological oscillations and response to external stimuli.

In this paper, we analytically derive the holding time distributions in the free and bound gene states for a full model of autoregulatory loops with both mRNA and protein descriptions. The paper is organized as follows. In Section 2, we describe the full model and discuss how it can be simplified to a reduced model with only the protein description when mRNA decays much faster compared to protein. In Section 3, we obtain the exact switching time distributions between the unbound and bound states for the full model under arbitrary initial conditions as well as under steady-state conditions; the analytical solution is found to be in excellent agreement with the numerical solution obtained using stochastic simulations. In Section 4, we show how the exact solution for the full model can be used to obtain the exact switching time distributions for the reduced model. In particular, we point out that the previous solution given in [26] for the reduced model is not exact due to the choice of an incorrect initial condition, and we further show how to correct this solution. In Section 5, we discuss the possible distribution shapes for the switching time distributions and examine the corresponding phase diagram. We find that the full model of an autoregulatory loop can produce three distinct shapes of steady-state holding time distributions: a unimodal distribution with a zero peak, a unimodal distribution with a nonzero peak, as well as a bimodal distribution with both a zero and a nonzero peak. The most interesting of these phases is a bimodal phase, and we show that this novel distribution type fails to be captured by the reduced model with only the protein description. We conclude in Section 6.

## 2 Model

### 2.1 Full model with both mRNA and protein descriptions

Here we consider a standard three-stage stochastic model of an autoregulatory feedback loop including transcription, translation, and protein-gene interactions (Fig. 1(a)) [10]. Due to feedback regulation, the gene can be either free or bound to a protein molecule. Let *G* and *G*^*^ denote the free (unbound) and bound states of the gene, respectively, let *M* denote the corresponding mRNA, and let *P* denote the corresponding protein. The autoregulatory loop can be described by the following reaction scheme:

**Figure 1.**
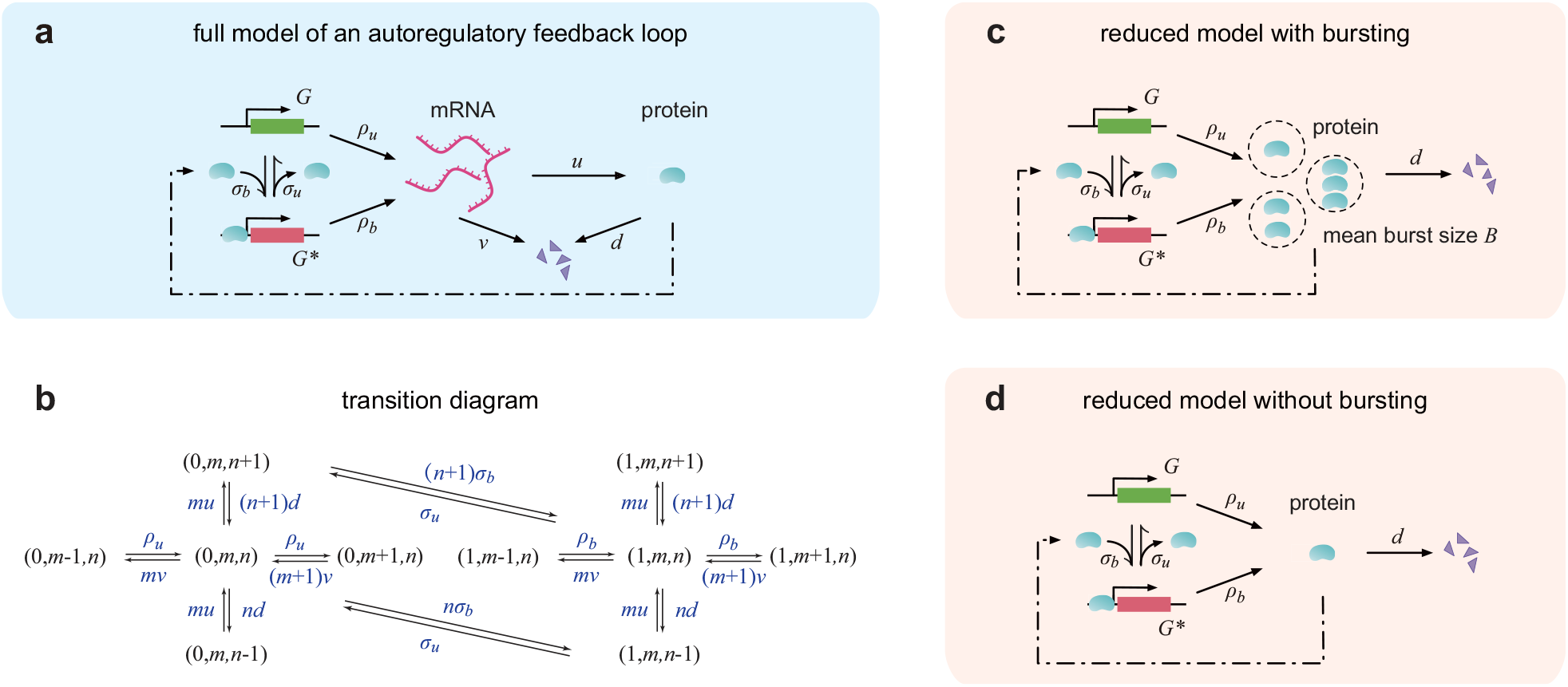
Models. **(a)** Full model of an autoregulatory feedback loop with transcription, translation, decay of mRNA and protein, and switching between an unbound and a bound gene state. Due to feedback regulation, the switching rate from the unbound to the bound state depends on the instantaneous protein copy number and hence the duration spent in the unbound gene state has a non-exponential distribution. **(b)** Transition diagram of the Markovian dynamics for the full model shown in (a). Here the microstate of the gene of interest is described by an ordered triple (*i, m, n*), where *i* denotes the gene state (*i* = 0 corresponds to the unbound state and *i* = 1 corresponds to the bound state), *m* denotes the mRNA number, and *n* denotes the protein number. **(c)** Reduced model with only the protein description. Here the mRNA dynamics is modelled implicitly. The reduced model can be viewed as the limit of the full model when mRNA decays much faster than protein (*v » d*). In this case, protein molecules are produced in a bursty manner. **(d)** Reduced model with only the protein description. Here protein molecules are produced in a non-bursty manner.

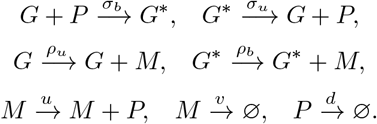

Here *σ*_*b*_ is the binding rate of protein to the gene; *σ*_*u*_ is the unbinding rate; *ρ*_*u*_ and *ρ*_*b*_ are the transcription rates in the unbound and bound gene states, respectively; *u* is the translation rate; *v* and *d* are the decay rates of mRNA and protein, respectively, either due to active degradation or due to dilution during cell division [27, 28]. The reaction system describes a positive feedback loop when ρ_*b*_ > *ρ*_*u*_ (binding of a protein promotes its own expression) and describes a negative feedback loop when *ρ*_*b*_ < *ρ*_*u*_ (binding of a protein inhibits its own expression).

The microstate of the gene of interest can be represented by an ordered triple (*i, m, n*), where *i* is the gene state with *i* = 0, 1 corresponding to the unbound and bound states, respectively, *m* is the mRNA copy number, and *n* is the protein copy number. The stochastic gene expression dynamics can be described by the Markov jump process illustrated in Fig. 1(b). Let *p*_*i*,*m*,*n*_ denote the probability of having *m* copies of mRNA and *n* copies of protein in a single cell when the gene is in state *i*. Then the evolution of the Markovian model is governed by the CMEs

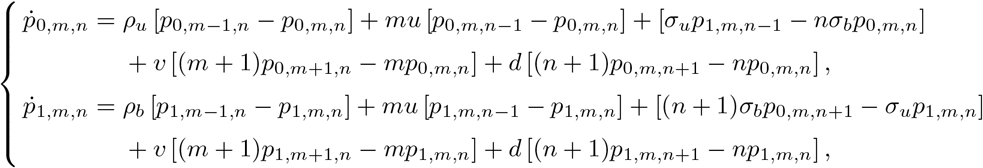

where the terms involving *ρ*_*u*_ and *ρ*_*b*_ on the right-hand side describe mRNA synthesis, the terms involving *u* describe protein synthesis, the terms involving *σ*_*u*_ and *σ*_*b*_ describe gene state switching and feedback regulation, the terms involving *v* describe mRNA decay, and the terms involving *d* describe protein decay.

### 2.2 Reduced model with only the protein description

In bacteria and yeast, mRNA often decays much faster than its protein counterpart [29]. In fact, mRNA lifetimes in prokaryote are generally on the order of a few minutes, while protein lifetimes are on the order of tens of minutes to many hours [30]. Let *γ* = *v/d* denote the ratio of the decay rates of mRNA and protein, and let *B* = *u/v* denote the average number of protein molecules produced per mRNA lifetime. Here we make the assumption that *γ »* 1 and *B* is strictly positive and bounded [10]. This guarantees that the process of protein synthesis followed by mRNA degradation is essentially instantaneous. Once an mRNA molecule is synthesized, it can either produce a protein molecule with probability *p* = *u/*(*u* + *v*) or be degraded with probability *q* = 1 − *p* = *v/*(*u* + *v*). The probability that a transcript produces *k* copies of protein before it is finally degraded will be *p*^*k*^*q*, which follows a geometric distribution. In this case, protein synthesis occurs in a bursty manner (it is widely known that bursty production of protein results from rapid translation of protein from a single, short-lived mRNA molecule [31, 32]). Since the protein burst size is geometrically distributed, its expected value is given by

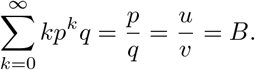

In this case, the full model with both mRNA and protein descriptions (Fig. 1(a)) can be reduced to a model with only the protein description (Fig. 1(c)). The reduced model can be described by the following reactions:

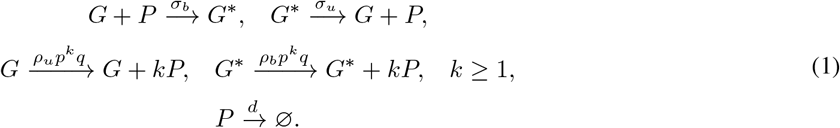

Note that in both the full and reduced models, the switching from the bound to the unbound gene state occurs at constant rate *σ*_*u*_, and thus the holding time in the bound gene state, i.e. the time spent in the bound state before switching to the unbound state, has an exponential distribution with probability density 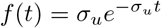. On the other hand, due to autoregulation, the switching from the unbound to the bound gene state occurs at a rate that depends on the instantaneous protein number *n* (Fig. 1(b)). Hence the holding time in the unbound gene state, i.e. the time spent in the unbound state before switching to the bound state, must have a non-exponential distribution. In this paper, we will compute the analytical distribution of the unbound duration, i.e. the holding time distribution in the unbound gene state, for both the full and reduced models of an autoregulatory circuit.

We emphasize that the analytical distribution of the unbound duration has been examined previously in [26] for a simpler model of autoregulatory circuits with only the protein description when protein synthesis occurs in a non-bursty manner (Fig. 1(d)). However, we find that the solution given in [26] is only approximate and is not exact (the detailed reasons will be explained later). Here we will generalize and correct the results obtained in previous studies by taking the mRNA dynamics (for the full model) and protein bursting (for the reduced model) into account.

## 3 Analytical unbound duration distributions for the full model

### 3.1 Analytical distributions under general initial conditions

We first derive the unbound duration distribution for the full model shown in Fig. 1(a). If we only focus on the evolution of the gene state, then we obtain a two-state stochastic trajectory that switches constantly between the unbound and bound states (Fig. 2(a)). For simplicity, we assume that the gene is in the unbound state at time *t* = 0. We next introduce some notation. Let 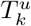 denote the *k*th *hitting time* of the unbound gene state, i.e. the time needed for the gene to enter the unbound state for the *k*th time. Similarly, let 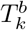 denote the *k*th hitting time of the bound gene state, i.e. the time needed for the gene to enter the bound state for the *k*th time. Note that both 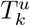 and 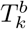 are random times (they are called stopping times in probability theory). Then the *k*th unbound duration, i.e. the *k*th switching time from the unbound to the bound state, is given by (see Fig. 2(a) for an illustration)

**Figure 2.**
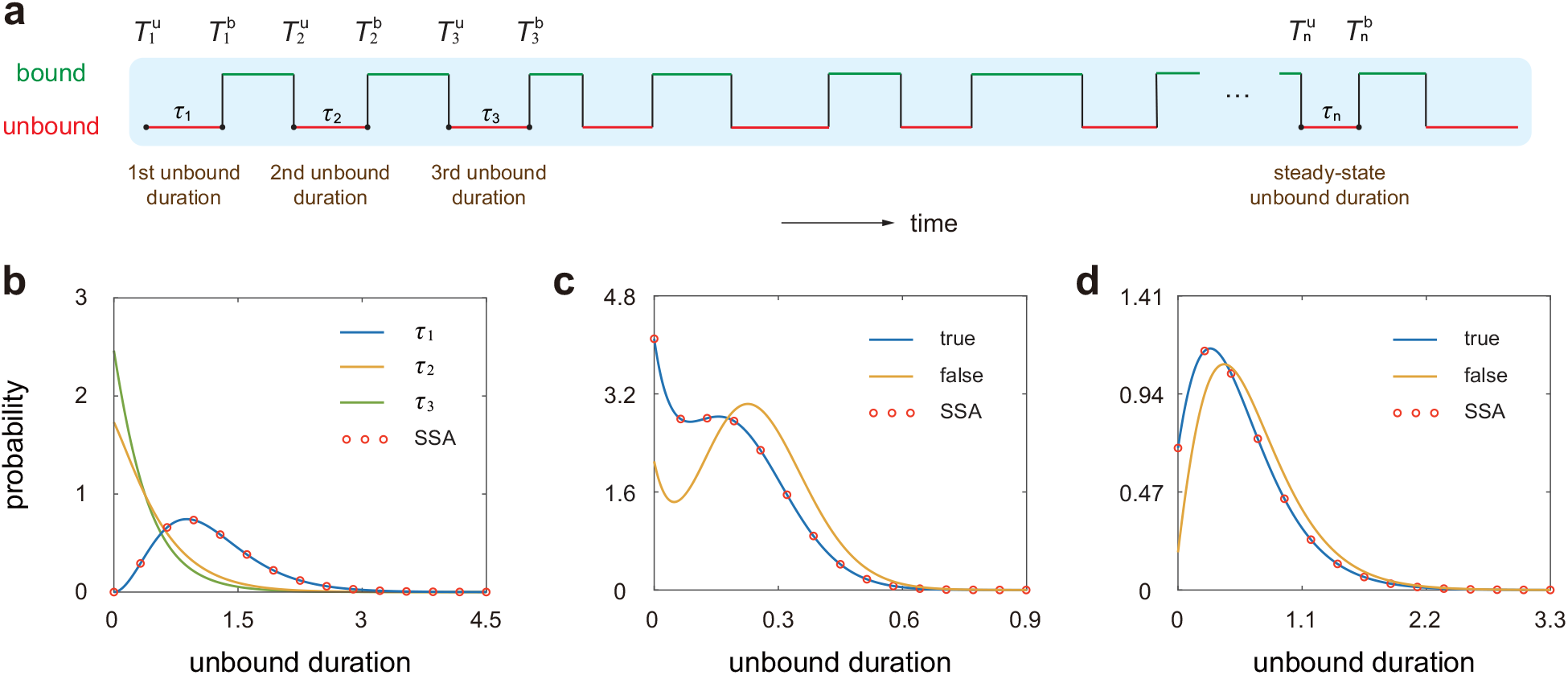
Unbound duration distributions in non-steady-state and steady-state conditions. **(a)** Schematics of a stochastic trajectory of the gene state in an individual cell. The gene switches between the unbound and bound states. The time spent in the unbound gene state is called the unbound duration (red lines). The distribution of the unbound duration after a sufficient number of switching events is called the steady-state unbound duration distribution. **(b)** Comparison between the distributions of the first (blue curve), second (yellow curve), and third (green curve) unbound durations when the gene is initially in the unbound state and when initially there are no mRNA and protein molecules in the cell. The blue curve shows the exact solution given in Eqs. (7) and (8), and the red circles show the simulated results obtained from the SSA. The parameters are chosen as *ρ*_*u*_ = 30, *ρ*_*b*_ = 60, *σ*_*u*_ = 10, *σ*_*b*_ = 0.1, *u* = *v* = 3, *d* = 1. **(c)** Comparison between the steady-state distributions of the unbound duration for the full model shown in Fig. 1(a) with the correct initial condition given in Eq. (9) (blue curve) and the incorrect one given in Eq. (12) (yellow curve). The red circles show the simulated results obtained using the SSA. The parameters are chosen as *ρ*_*u*_ = 60, *ρ*_*b*_ = 0, *σ*_*u*_ = 0.1, *σ*_*b*_ = 2, *u* = *v* = 3, *d* = 1. **(d)** Comparison between the steady-state distributions of the unbound duration for the reduced model shown in Fig. 1(d) with the correct initial condition given in Eq. (18) (blue curve) and the incorrect one given in Eq. (19) (yellow curve). The red circles show the simulated results obtained using the SSA. The parameters are chosen as *ρ*_*u*_ = 10, *ρ*_*b*_ = 0, *σ*_*u*_ = 0.1, *σ*_*b*_ = 0.1, *d* = 1.

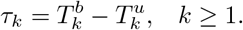

we now compute the distribution of the first unbound duration *τ*_1_, i.e. the first period of time spent in the unbound state, under arbitrary initial conditions. Note that *τ*_1_ is also referred to as the first passage time to switch from the unbound to the bound gene state in previous studies [26]. To this end, we denote by *F* (*t*) = ℙ (*τ*_1_ *> t*) the survival function of *τ*_1_, i.e. the probability that the gene remains in the unbound state during the time interval [0, *t*]. Once we have obtained the explicit expression of *F* (*t*), the probability density of *τ*_1_ can be obtained by taking the negative derivative of *F* (*t*).

To proceed, let *h*(*m, n, t*) denote the probability of having *m* copies of mRNA and *n* copies of protein in a single cell when the gene remains in the unbound state during the time interval [0, *t*]. It is clear that

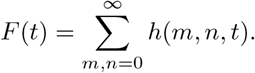

we now derive the evolution equation satisfied by *h*(*m, n, t*). Suppose that there are *m* copies of mRNA and *n* copies of protein in a single cell at time *t* and the gene remains in the unbound state during the time interval [0, *t*]. Since the gene state does not change during [0, *t*], within the small time interval [*t* − ∆*t, t*], only four reactions can occur and the corresponding changes in microstate are given as follows:

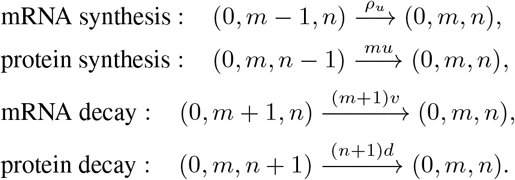

The probability for each reaction to occur during [*t* − ∆*t, t*] is approximately the propensity of the reaction multiplied by the length of the interval. For example, the probability that a transcript is synthesized during [*t* − ∆*t, t*] is *ρ*_*u*_∆*t* + *o*(∆*t*) and the probability that a protein molecule is synthesized is *mu*∆*t* + *o*(∆*t*). On the other hand, it is clear that the probability that no reactions occur during [*t* − ∆*t, t*] is 1 − (*ρ*_*u*_ + *mu* + *mv* + *nd* + *nσ*_*b*_) ∆*t* + *o*(∆*t*). Hence it follows from Bayes’ rule that

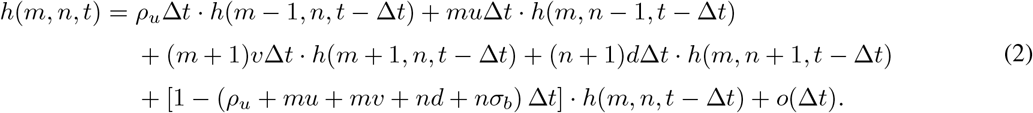

Subtracting *h*(*m, n, t* − ∆*t*) on both sides of Eq. (2) and letting ∆*t* → 0, we find that *h*(*m, n, t*) is governed by the following ordinary differential equations (ODEs):

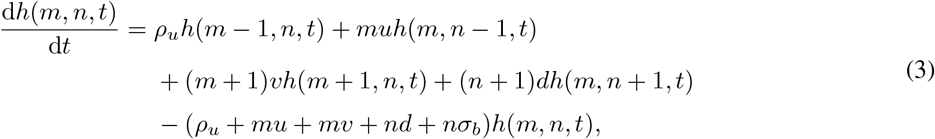

together with the initial condition *h*(*m, n*, 0) = *p*_0,*m*,*n*_(0), where *p*_0,*m*,*n*_(0) is probability of the system being in microstate (0, *m, n*) at time *t* = 0.

To solve Eq. (3), we define the generating function

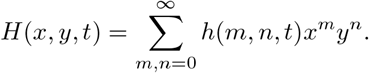

Then Eq. (3) can be converted into the following first-order linear partial differential equation (PDE):

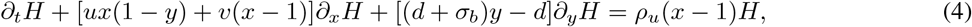

together with the initial condition

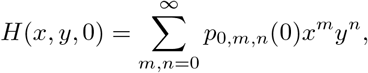

which can be determined by the initial distribution of the system. Using the method of characteristics, the generating function *H*(*x, y, t*) can be computed analytically as (see Appendix A for the proof)

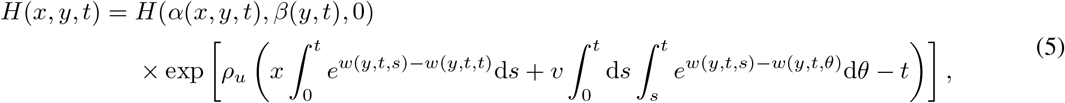

where

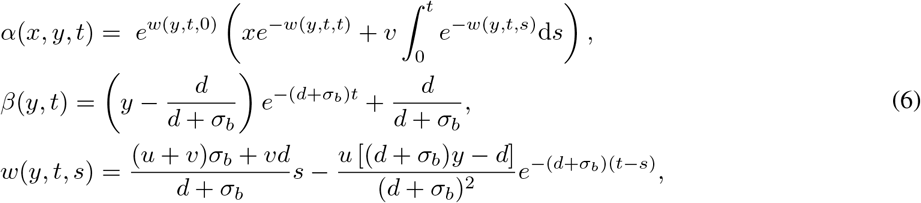

are three functions. Then the survival function *F* (*t*) of the unbound duration *τ*_1_ can be recovered from the generating function *H*(*x, y, t*) by taking *x* = *y* = 1, i.e.

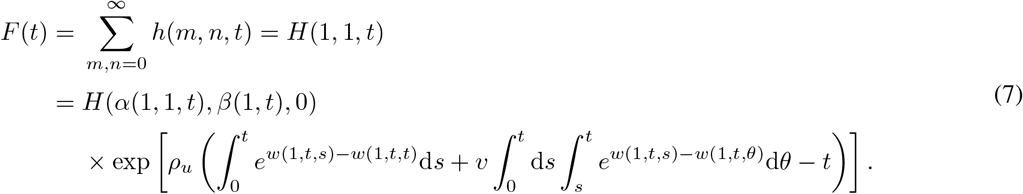

Finally, the probability density of the unbound duration *τ*_1_ is given by

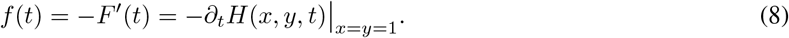

Thus far, we have obtained the analytical distribution of the unbound period *τ*_1_ under arbitrary initial conditions for the full model of an autoregulatory circuit. Similarly, we can compute the exact distribution of the *k*th unbound period *τ*_*k*_ for any *k* ≥ 2 (see Appendix B for details). Note that the distribution of *τ*_*k*_ is simply the distribution of *τ*_1_ with the initial distribution of mRNA and protein numbers, i.e. *p*_0,*m*,*n*_(0), replaced by the mRNA and protein number distribution at the hitting time 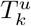 (these distributions are referred to as *hitting distributions* in probability theory). To validate our analytical solution, we compare it with the numerical solution obtained using the stochastic simulation algorithm (SSA) under the condition that the gene is initially in the unbound state and initially there are no mRNA and protein molecules in the cell (Fig. 2(b)). This mimics the situation where the gene has been silenced by some repressor over a period of time such that all mRNA and protein molecules have been removed via degradation. At time *t* = 0, the repressor is removed and we study how gene expression recovers. Clearly, the two solutions are in perfect agreement.

### 3.2 Analytical distributions in steady-state conditions

We have obtained the exact distribution of the unbound period under arbitrary initial conditions. From Fig. 2(b), it can be seen that the distributions of *τ*_1_, *τ*_2_, and *τ*_3_ are generally different. However, after a sufficient number of switching events, the system will reach a steady state and the distribution of *τ*_*k*_ will become independent of *k*. We next derive the analytical distribution of the unbound duration in steady-state conditions, i.e. the holding time distribution in the unbound gene state after a sufficient number of switching events.

Note that after a large number of switching events, the joint distribution of mRNA and protein numbers at the hitting time 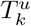 will also become independent of *k* and this distribution will be called the *steady-state hitting distribution* in the following. We make a crucial observation that the steady-state distribution of the unbound duration is exactly the distribution of the first unbound duration *τ*_1_ with the initial mRNA and protein number distribution replaced by the steady-state hitting distribution (Fig. 2(a)).

We next compute the steady-state hitting distribution. To this end, let

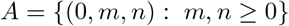

be the collection of all microstates in the unbound gene state and let

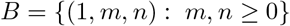

be the collection of all microstates in the bound gene state. In Appendix B, we prove that for any microstate (0, *m, n*) in the set *A*, the steady-state hitting probability of this microstate is exactly the total steady-state probability flux from the set *B* to this microstate divided by the total steady-state probability flux from the set *B* to the set *A*. From Fig. 1(b), it is clear that the transition from microstate (1, *m, n* − 1) to microstate (0, *m, n*) (with rate *σ*_*u*_) is the only possible transition from the set *B* to microstate (0, *m, n*). Hence the steady-state hitting distribution is given by

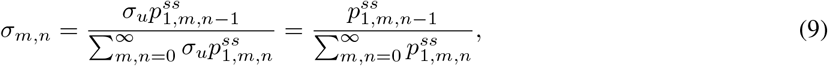

where 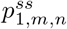 is the steady-state probability of the system being in microstate (1, *m, n*), 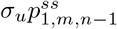 is the steady-state probability flux from the set *B* to microstate (0, *m, n*), and 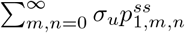 is the total steady-state probability flux from the set *B* to the set *A*. Finally, we obtain the exact distribution of the unbound duration in steady-state conditions, which is still given by Eqs. (7) and (8), with the initial condition *H*(*x, y*, 0) replaced by

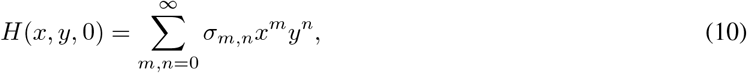

where *σ*_*m*,*n*_ is the steady-state hitting distribution given in Eq. (9).

Note that in order to analytically compute the steady-state unbound duration distribution, we need to first obtain the steady-state joint distribution of mRNA and protein numbers since *σ*_*m*,*n*_ depends on 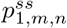 . We emphasize that thus far, the steady-state mRNA and protein number distribution for the full model shown in Fig. 1(a) has not been obtained analytically. However, it can be solved in closed form in some special cases, such as in the case of weak feedback [33]. In Appendix C, we derive the exact steady-state joint distribution of mRNA and protein numbers when the gene switches very slowly between the two states, i.e. *σ*_*u*_, *σ*_*b*_ ≪ *ρ*_*u*_, *ρ*_*b*_, *u, v, d*. In this limiting case, the analytical solution is given by

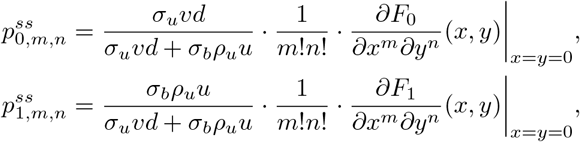

where

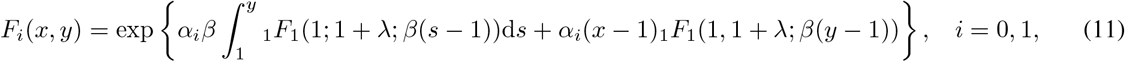

with _1_*F*_1_(*a*; *b*; *z*) being Kummer’s confluent hypergeometric function [34], and with

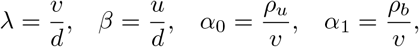

being four constants. In the general case, we may use the finite-state projection (FSP) algorithm [35] to compute the steady-state mRNA and protein number distribution, as well as the steady-state hitting distribution *σ*_*m*,*n*_.

We emphasize that in previous studies [26], the steady-state distribution of the unbound duration has been obtained in a simpler model of autoregulatory loops with only the protein description. In that study, the steady-state hitting distribution *σ*_*m*,*n*_ was incorrectly chosen as the steady-state conditional distribution in the bound gene state, i.e.

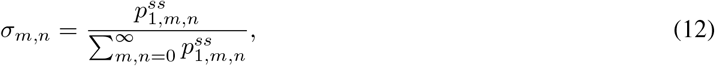

since the authors believed that “it was the distribution seen by the system when the reaction *G* + *P* → *G*^*^ became first possible” [26]. Comparing Eqs. (9) and (12), it can be seen that the initial condition used in previous papers is

similar to the correct one but is still not exact. Fig. 2(c) compares the steady-state unbound period distributions with the correct initial condition given by Eq. (9) and with the incorrect one given by Eq. (12). Clearly, the two distributions may deviate significantly from each other.

### 3.3 Numerical validation and heavy-tailed distributions

To test our analytical solution given by Eqs. (7) and (8) with the initial condition given by Eq. (10), we compare it with the numerical solution obtained from the SSA for both positive and negative feedback loops (Fig. 3). When performing the SSA, we compute the steady-state distribution of unbound durations by generating a single stochastic trajectory with 10^4^ switching events from the unbound to the bound gene state. As expected, the two solutions coincide excellently. We find that when the binding and unbinding rates are not very large, the analytical solution is computationally much more efficient than the SSA. This is because when gene switching is relatively slow, the majority of the time in the SSA is spent simulating synthesis and decay of the gene products, and we need to generate a very long trajectory in order to obtain enough gene switching events. According to simulations, the full model of an autoregulatory loop with both mRNA and protein descriptions can produce three distinct shapes of steady-state unbound period distributions: a unimodal distribution with a zero peak (decaying distribution), a unimodal distribution with a nonzero peak (bell-shaped distribution), and a bimodal distribution with both a zero and a nonzero peak. Interestingly, we find that a negative feedback loop can exhibit all the three shapes of distributions (Fig. 3(a)), while a positive feedback loop can only display a decaying or a bell-shaped distribution (Fig. 3(b)). We emphasize that in previous studies [25, 26], it was found that the unbound period may possess a bell-shaped distribution in a negative feedback model with only the protein description. Here we find that if the mRNA dynamics is modelled explicitly, then a negative feedback loop may even exhibit a bimodal distribution.

**Figure 3.**
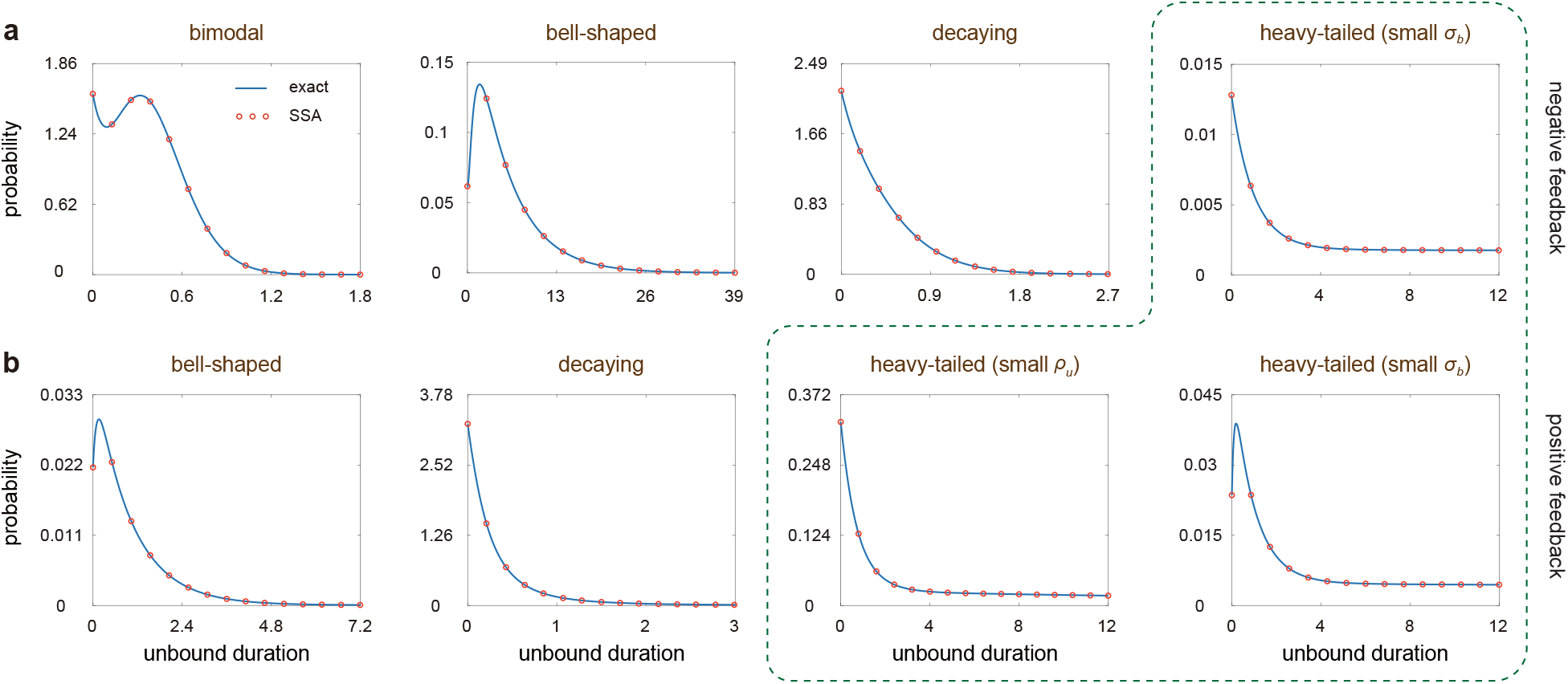
Different shapes of steady-state unbound duration distributions for the full model of an autoregulatory feedback loop. **(a)** A negative feedback loop can display a decaying, a bell-shaped, or a bimodal distribution. The parameters are chosen as *ρ*_*u*_ = 30, *ρ*_*b*_ = 0, *σ*_*u*_ = 0.1, *σ*_*b*_ = 0.8, *u* = *v* = 3, *d* = 1 (first column), *ρ*_*u*_ = 20, *ρ*_*b*_ = 5, *σ*_*u*_ = 0.01, *σ*_*b*_ = 0.01, *u* = *v* = 5, *d* = 1 (second column), *ρ*_*u*_ = 40, *ρ*_*b*_ = 20, *σ*_*u*_ = 0.1, *σ*_*b*_ = 0.1, *u* = *v* = 5, *d* = 1 (third column), and *ρ*_*u*_ = 10, *ρ*_*b*_ = 0, *σ*_*u*_ = 10, *σ*_*b*_ = 0.011, *u* = 0.1, *v* = 6, *d* = 1 (fourth column). **(b)** A positive feedback loop can only display a decaying or a bell-shaped distribution. The parameters are chosen as *ρ*_*u*_ = 0.02, *ρ*_*b*_ = 50, *σ*_*u*_ = 10, *σ*_*b*_ = 0.002, *u* = 30, *v* = 10, *d* = 1 (first column), *ρ*_*u*_ = 5, *ρ*_*b*_ = 20, *σ*_*u*_ = 0.16, *σ*_*b*_ = 0.16, *u* = *v* = 10, *d* = 1 (second column), *ρ*_*u*_ = 0.1, *ρ*_*b*_ = 0.2, *σ*_*u*_ = 0.01, *σ*_*b*_ = 0.2, *u* = 9, *v* = 3, *d* = 1 (third column), and *ρ*_*u*_ = 0.5, *ρ*_*b*_ = 50, *σ*_*u*_ = 10, *σ*_*b*_ = 0.002, *u* = 20, *v* = 10, *d* = 1 (fourth column). In (a), (b), the blue curves show the analytical solution given in Eqs. (7) and (8) with the initial condition given by Eq. (10), and the red circles show the simulated results obtained from the SSA. The green dashed curve encloses heavy-tailed distributions, which generally occur when *σ*_*b*_ or *ρ*_*u*_ is small.

Furthermore, we also observe that in autoregulatory circuits, the steady-state unbound duration distribution may decay very fast with a short tail (Fig. 3, distributions outside the green dashed curve) or decay very slowly with a long tail (Fig. 3, distributions inside the green dashed curve). According to simulations, a heavy-tailed distribution is more likely to occur when the binding rate *σ*_*b*_ and the transcription rate in the unbound gene state *ρ*_*u*_ are very small compared to other rate constants. The heavy-tailed holding time distribution indicates the presence of long-lived switching events with a small probability within a population of cells, which may play an important role in driving slow dynamics of translational bursting. A more detailed explanation of the heavy-tailed distribution will be given later.

## 4 Analytical unbound duration distributions for the reduced model

### 4.1 Model with translational bursting

We next derive the exact distribution of the unbound duration for the reduced model shown in Fig. 1(c), whose reaction scheme is given by Eq. (1). Here the mRNA dynamics is modelled implicitly and protein molecules are produced in a bursty manner. For the reduced model, the microstate of the gene of interest can be represented by an ordered pair (*i, n*), where *i* = 0, 1 is the gene state and *n* is the protein number. Similarly, let *h*(*n, t*) denote the probability of having *n* copies of protein in a single cell at time *t* when the gene remains in the unbound state during the time interval [0, *t*]. Moreover, let

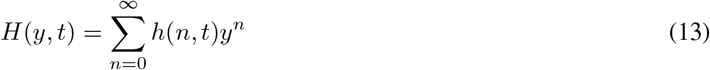

denote the corresponding generating function. For simplicity, we only focus on the unbound period distribution in steady-state conditions.

Recall that the reduced model can be viewed as the limit of the full model shown in Fig. 1(a) when mRNA decays much faster compared to protein (*γ* = *v/d* ≫ 1) and when the number of protein molecules produced per mRNA lifetime (*B* = *u/v*) is strictly positive and bounded [32, 36]. Taking the limit of *γ* → ∞ while keeping *u/v* as constant in Eq. (5), we find that the generating function *H*(*y, t*) can be computed explicitly as (see Supplementary Section 1 for details)

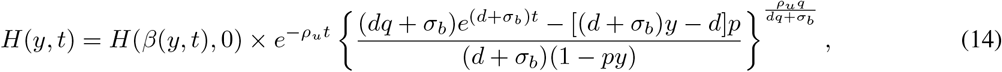

where *β*(*y, t*) is the function defined in Eq. (6). The initial condition of the generating function is determined by

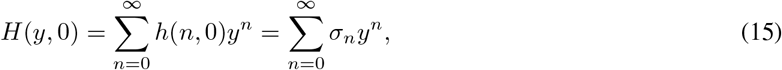

where

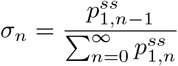

is the steady-state hitting distribution, i.e. the protein number distribution just after switching from the bound to the unbound gene state after a sufficient number of switching events. Here 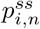 denotes the steady-state probability of having *n* protein copies when the gene is in state *i*. Its analytical solution has been obtained in [10] and is given by

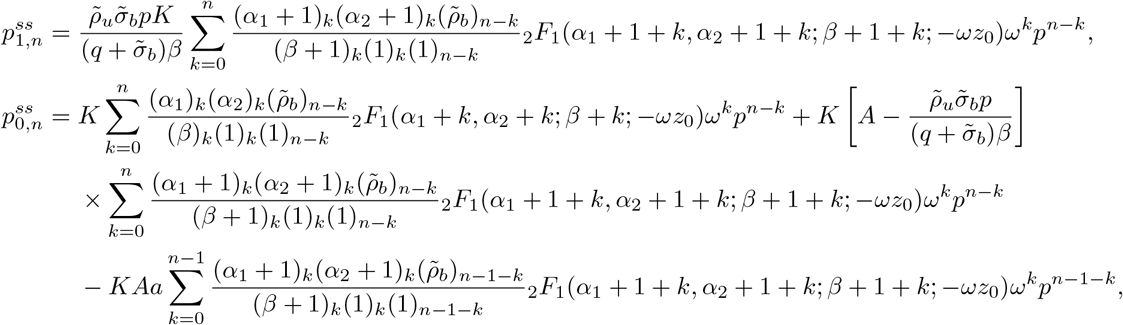

where (*x*)_*n*_ = *x*(*x* + 1) · · · (*x* + *n* − 1) denotes the Pochhammer symbol, _2_*F*_1_(*α*_1_, *α*_2_; *β*; *z*) denotes the Gaussian hypergeometric function [34], and

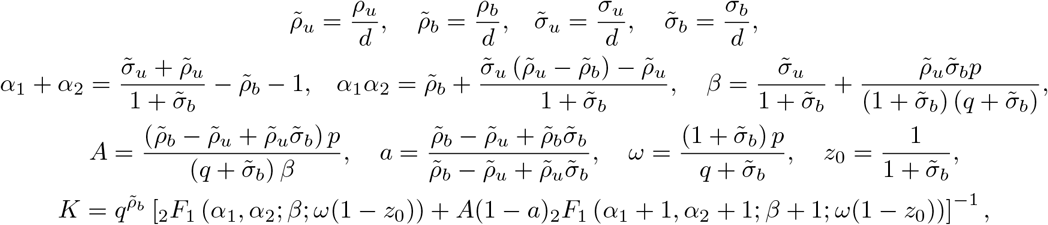

are some constants. Finally, the steady-state probability density of the unbound duration can be recovered by taking the negative derivative of *H*(1, *t*) with respect of time *t*, i.e.

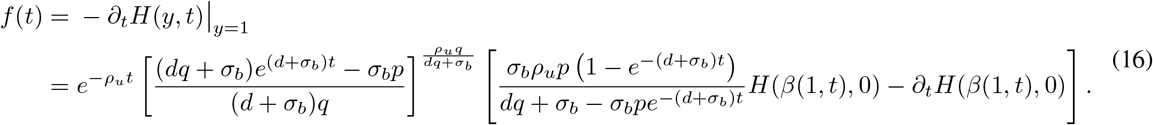

Recall that an autoregulatory loop can display a heavy-tailed distribution of the unbound duration (Fig. 3). This can be explained using the exact solution given in Eq. (16). When *t* ≫ 1, the term in the first bracket in Eq. (16) can be approximated by

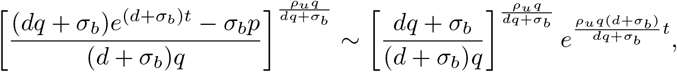

and the term in the second bracket in Eq. (16) is approximately a constant. Hence for large *t*, the unbound duration distribution can be approximated by

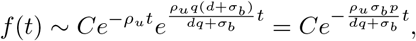

with *C >* 0 being a constant, and hence will decay exponentially with rate *ρ*_*u*_*σ*_*b*_*p/*(*dq* + *σ*_*b*_). Note that when the binding rate *σ*_*b*_ is small or the transcription rate in the unbound state *ρ*_*u*_ is small (note that a small *ρ*_*u*_ can only occur in positive feedback loops; see Fig. 3(b)), the exponential decay rate will be very close to zero and hence the unbound duration distribution will have a very long tail. This explains our previous finding that heavy-tailed distributions tend to occur when *σ*_*b*_ and *ρ*_*u*_ are small.

### 4.2 Model without translational bursting

Note that in the reduced model shown in Fig. 1(c), protein synthesis occurs in a bursty manner. In previous studies, a simpler reduced model with non-bursty protein synthesis has been considered [13, 26] and the unbound duration distribution has been computed analytically for that model [26]. If the mRNA lifetime is long (as commonly observed in mammalian cells [37]), then protein synthesis may appear to be non-bursty. The non-bursty reduced model of an autoregulatory loop is described by the following reactions (see Fig. 1(d) for an illustration):

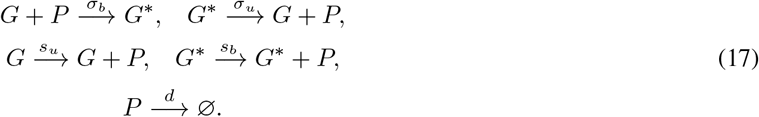

Here protein molecules are produced one at a time with rate *s*_*u*_ (*s*_*b*_) when the gene is in the unbound (bound) state.

In fact, the non-bursty reduced model given by Eq. (17) can be viewed as a limiting case of the bursty reduced model given by Eq. (1) [15, 38]. Note that when *ρ*_*u*_, *ρ*_*b*_ → ∞ and *p* → 0, while keeping *ρ*_*u*_*p* = *s*_*u*_ and *ρ*_*b*_*p* = *s*_*b*_ as constant, we have

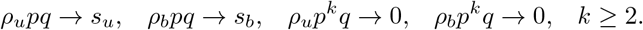

This shows that the bursty model reduces to the non-bursty one in the above limit. Hence the exact unbound duration distribution for the non-bursty model can be derived from that for the bursty model (Eqs. (14) and (16)) by taking the above limit. In this way, we can obtain the generation function *H*(*y, t*) for the non-bursty model, which is given by

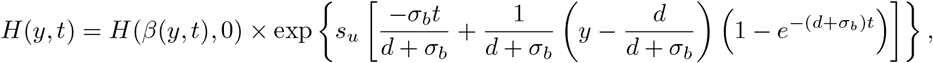

where *β*(*y, t*) is the function defined in Eq. (6). The initial condition *H*(*y*, 0) of the generating function is still determined by Eq. (15) with

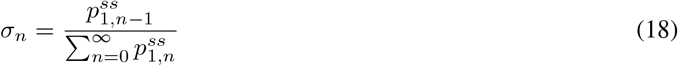

being the steady-state hitting distribution. Here 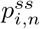 denotes the steady-state probability of having *n* protein copies when the gene is in state *i*. Its analytical solution has been obtained in [10] and is given by

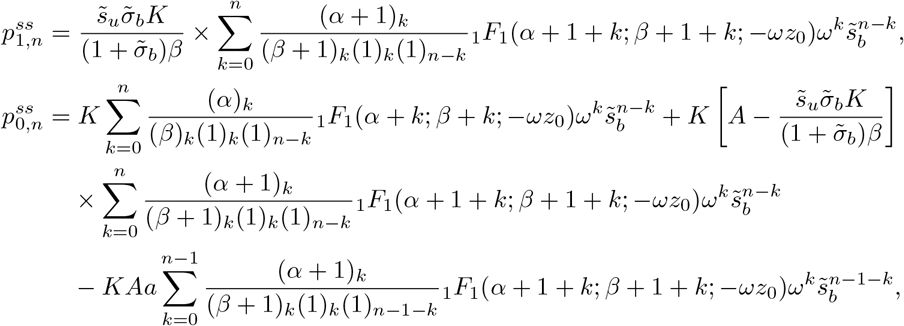

where _1_*F*_1_(*α*; *β*; *z*) is Kummer’s confluent hypergeometric function [34], and

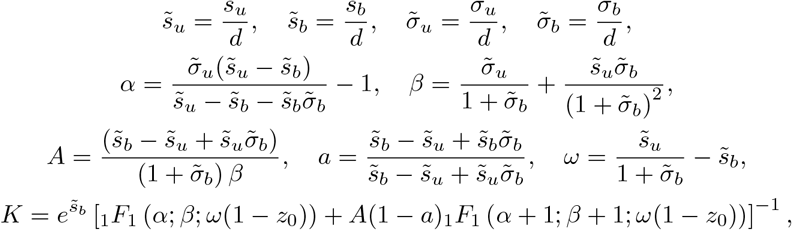

are some constants. Finally, the steady-state probability density of the unbound duration can be recovered by taking the negative derivative of *H*(1, *t*) with respect of time *t*, i.e.

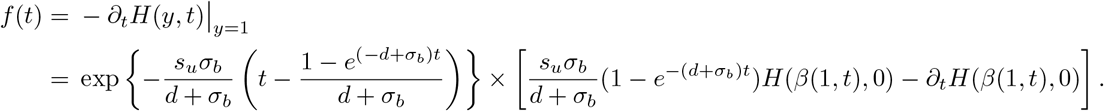

we emphasize that in previous studies [26], the steady-state unbound period distribution has been derived for the non-bursty reduced model; however, the steady-state hitting distribution *σ*_*n*_ was incorrectly chosen to be the conditional distribution of the protein number in the bound gene state, i.e.

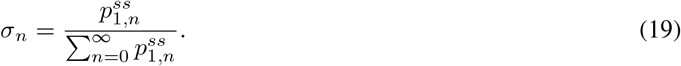

Fig. 2(d) compares the steady-state analytical distributions of the unbound duration computed using the correct initial condition given by Eq. (18) and the incorrect one given by Eq. (19). Clearly, there may be a significant difference between them, even when the mRNA dynamics is ignored.

## 5 Phase diagram of distribution shapes

We next examine the shape of the steady-state unbound duration distribution for the full model with both mRNA and protein descriptions (Fig. 1(a)) and the reduced model with only the protein description and with translational bursting (Fig. 1(c)). Recall that the latter can be viewed as the limit of the former when mRNA decays much faster compared to protein (*v* ≫ *d*). Our analytical solution and numerical solution obtained from the SSA indicate that when model parameters are appropriately chosen, the full model is capable of producing three shapes of distributions: a decaying distribution, a bell-shaped distribution, and a bimodal distribution (Fig. 3).

To determine the regions for the three distribution shapes in the parameter space, we illustrate the *σ*_*b*_ - *σ*_*u*_ phase diagrams for both the full and reduced models (Fig. 4). We first focus on negative feedback loops (Fig. 4(a)), which are capable of producing all the three distribution shapes. Interestingly, we find that the shape of the unbound duration distribution is closely related to the binding and unbinding rates. In the regime of slow gene switching (small *σ*_*b*_, *σ*_*u*_), the unbound duration generally has a bell-shaped distribution; in the regime of fast gene switching (large *σ*_*b*_, *σ*_*u*_), the unbound duration generally has a decaying distribution. The bimodal distribution occurs when the binding and unbinding rates are neither too large nor too small. As the mRNA decay rate *v* increases, the bell-shaped parameter region enlarges, while the bimodal and decaying parameter regions shrink simultaneously. In the limit of *γ* → ∞, the bimodal region totally disappears, which indicates that the reduced model can only produce decaying and bell-shaped distributions. This is consistent with the finding in previous studies [25, 26].

**Figure 4.**
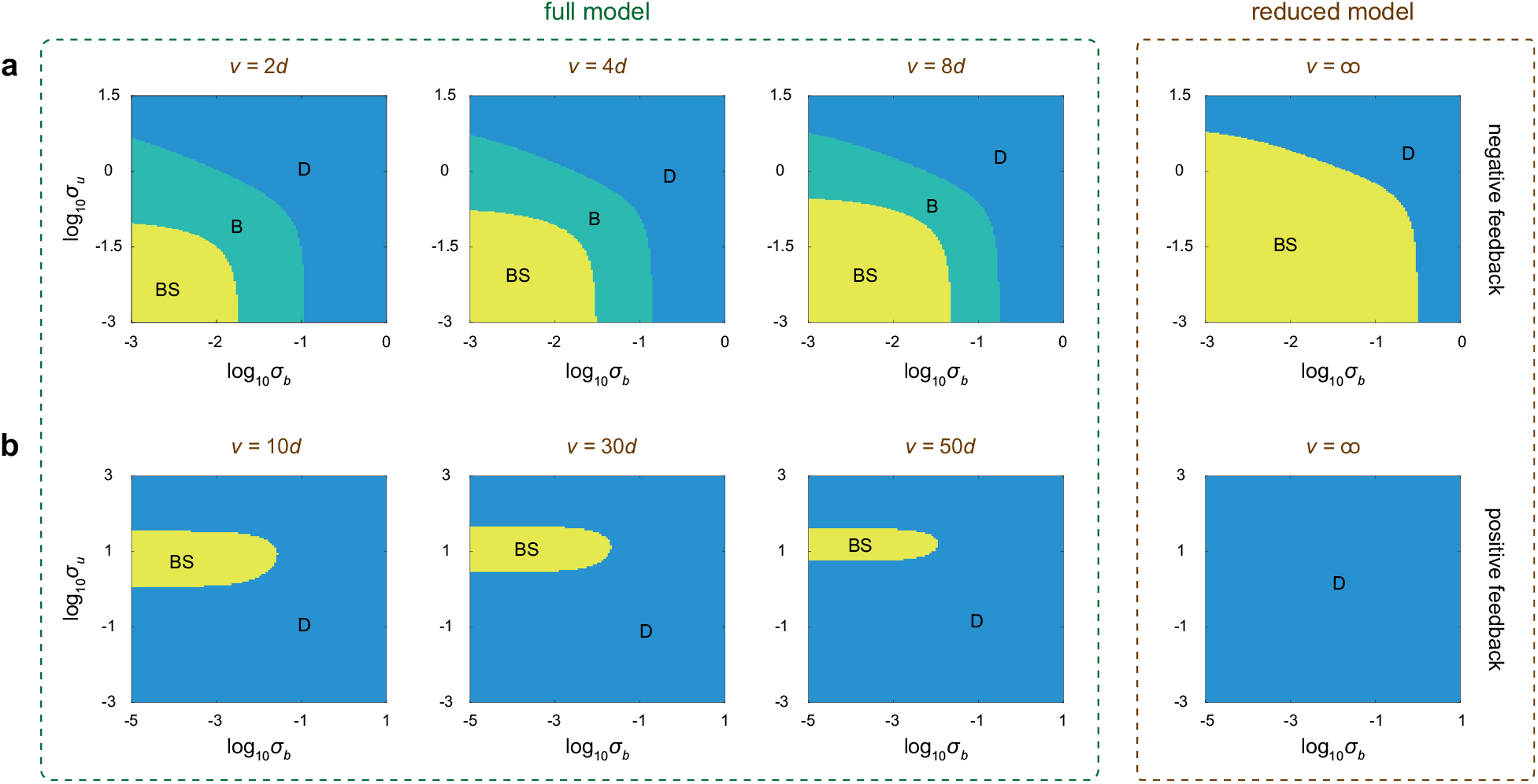
Phase diagrams of distribution shapes in the *σ*_*b*_ - *σ*_*u*_ plane for the full and reduced models of positive and negative feedback loops. **(a)** Phase diagrams of negative feedback systems. The system can produce a decaying (D), a bell-shaped (BS), or a bimodal (B) steady-state distribution of the unbound duration. The parameters are chosen as *ρ*_*u*_ = 20, *ρ*_*b*_ = 5, *d* = 1, *u* = *v*. The decay rate of mRNA is chosen as *v* = 2*d, v* = 4*d, v* = 8*d*, and *v* = ∞. Note where *v* =∞ corresponds to the reduced model since the reduced model can be viewed as the limit of the full model when *γ* = *v/d*→ ∞ and when *u/v* remains strictly positive and bounded. **(b)** Phase diagrams of positive feedback systems. The system can produce a decaying (D) or a bell-shaped (BS) steady-state distribution. The parameters are chosen as *ρ*_*u*_ = 5, *ρ*_*b*_ = 20, *d* = 1, *u* = *v*. The decay rate of mRNA is chosen as *v* = 10*d, v* = 30*d, v* = 50*d*, and *v* = ∞. Again, *v* = ∞ corresponds to the reduced model.

The situation is totally different in positive feedback loops (Fig. 4(b)). Unlike negative feedback systems, a positive feedback loop can only display a decaying or a bell-shaped distribution of the unbound duration. A bell-shaped distribution can only take place when the binding rate *σ*_*b*_ is small and when the unbinding rate *σ*_*u*_ is controlled within a narrow range. In particular, a bell-shaped distribution fails to be observed when the gene switches very slow or very fast between the two states, or when the binding rate is relatively large. With the mRNA decay rate *v* increases, the bell-shaped parameter region shrinks. In the limit of *γ* → ∞, the bell-shaped region totally disappears, which indicates that the reduced model can only exhibit a decaying distribution.

In summary, we find that the full model of a negative feedback loop can display three shapes of unbound duration distributions, while the reduced model fails to exhibit a bimodal distribution. Similarly, the full model of a positive feedback loop can display a decaying or a bell-shaped distribution, while the reduced model fails to exhibit a bell-shaped distribution. This suggests that negative feedback systems are easier to produce complex distribution shapes than positive feedback ones. In both the positive and negative feedback cases, the unbound duration has a decaying distribution in the fast switching regime that is close to an exponential one. Note that even when mRNA decays much faster than protein, which is common in most cell types such as in bacteria, yeast, and mammalian cells [29] (in mouse fibroblasts, the median mRNA half-life is 9 h and the median protein half-life is 46 h [37]; in this case we have *v* ≈ 5*d*), the full model can still produce a distribution shape that the reduced model fails to capture (Fig. 4). This demonstrates that a model that ignores the mRNA dynamics does not always capture the realistic distribution shape of the unbound duration in steady-state conditions.

To further understand the distribution shape of the unbound duration, we depict the *ρ*_*u*_ - *ρ*_*b*_ phase diagrams for both the full and reduced models when the binding and unbinding rates are relatively small (Fig. 5(a)). Note that in the case, a positive feedback loop (*ρ*_*u*_ < *ρ*_*b*_) can only exhibit a decaying distribution (Fig. 4(b)), while a negative feedback loop (*ρ*_*u*_ > *ρ*_*b*_) can produce all the three shapes of distributions, depending on the size of *ρ*_*u*_ and *ρ*_*b*_. As the mRNA decay rate *v* increases, the bimodal parameter region shrinks. Again, in the limit of *γ* → ∞, the bimodal region totally disappears, which indicates that the reduced model of a negative feedback system can only exhibit a decaying or a bell-shaped distribution. When *ρ*_*b*_ is fixed while increasing *ρ*_*u*_, we find that the system undergoes a triphasic stochastic bifurcation from the decaying phase to the bimodal phase, and then to the bell-shaped phase (Fig. 5(b)). Hence in negative feedback loops, a bell-shaped unbound duration distribution is more likely to occur when the transcription rates in two gene states differ significantly (*ρ*_*u*_ ≫ *ρ*_*b*_), while a bimodal distribution tends to occur when *ρ*_*u*_ is neither too small nor too large compared to *ρ*_*b*_.

**Figure 5.**
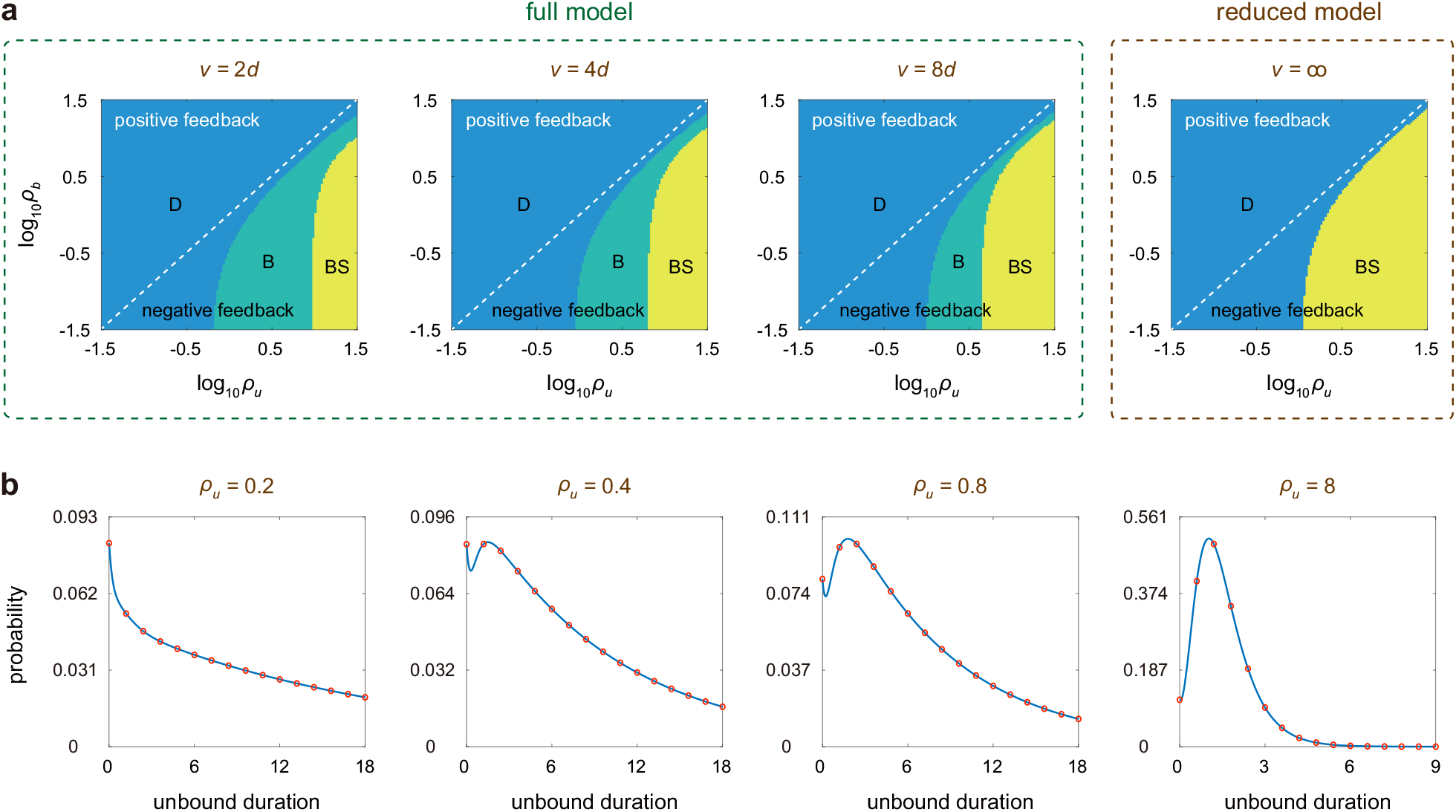
Effects of the transcription rates on the shape of the steady-state unbound duration distribution. **(a)** Phase diagrams of distribution shapes in the *ρ*_*u*_ - *ρ*_*b*_ plane. The white dashed line separates the parameter regions of positive and negative feedback systems. The parameters are chosen as *σ*_*u*_ = *σ*_*b*_ = 0.01, *d* = 1, *u* = *v*. The decay rate of mRNA is chosen as *v* = 2*d, v* = 4*d, v* = 8*d*, and *v* =*∞* . Note where *v* = *∞* corresponds to the reduced model. **(b)** Steady-state unbound period distributions under different choices of *ρu*. As *ρu* increases, the system undergoes a triphasic stochastic bifurcation from the decaying phase to the bimodal phase, and then to the bell-shaped phase. The parameters are chosen as *ρ*_*b*_ = 0, *σ*_*u*_ = 0.03, *σ*_*b*_ = 0.07, *u* = 9, *v* = 3, *d* = 1.

Note that compared to a decaying distribution with the same mean, a bell-shaped distribution generally has smaller noise. From Fig. 5(a), a negative feedback system is more likely to produce a bell-shaped unbound duration distribution than a positive feedback one. This suggests that negative feedback loops are more capable of producing an unbound duration with small noise. This “approximately deterministic” unbound duration may result in periodic switching from the free to the bound state and thus may further drive robust biological oscillations. This also reinforces previous findings that sustained oscillations are more likely to be observed in a negative feedback loop [39, 40].

## 6 Conclusions and discussion

In this work, we analytically derived the switching time distributions between the free and bound gene states, both under arbitrary initial conditions and under steady-state conditions, for (i) the full model of an autoregulatory feedback loop with both mRNA and protein descriptions and (ii) the reduced model with only the protein description and with translational bursting. The full model includes transcription, translation, and protein-gene interactions; the reduced model can be viewed as the limit of the full model when mRNA decays much faster than protein. Our exact solution generalizes and corrects previous ones in two different aspects: (i) in previous studies [25, 26], the switching time distributions have only been obtained in closed form in a simple model of an autoregulatory loop when the mRNA dynamics is ignored and when protein molecules are produced in a non-bursty manner. Here we generalized previous results by taking the mRNA dynamics (for the full model) and translational bursting (for the reduced model) into account; (ii) in previous studies [25, 26], the initial condition for the steady-state solution, i.e. the mRNA and protein number distribution just after switching from the bound to the unbound state, was chosen as the conditional steady-state distribution in the bound gene state. Here we showed that this is not exact, and the corrected one should be the steady-state hitting distribution, i.e. the mRNA and protein number distribution just after switching from the bound to the unbound state after a sufficient number of switching events.

Previous studies [25, 26] have shown that for a simple autoregulatory model with only the protein description, the holding time in the unbound gene state can only display a unimodal distribution with either a zero peak (decaying distribution) or a nonzero peak (bell-shaped distribution). Interestingly, here we showed that if the mRNA dynamics is taken into account, then the holding time in the unbound state can even exhibit a bimodal distribution with both a zero and a nonzero peak. Specifically, the full model of a negative feedback loop can display all the three shapes of distributions, while the reduced model can never produce a bimodal distribution; moreover, the full model of a positive feedback loop can display a decaying or a bell-shaped distribution, while the reduced model can only produce a decaying distribution. This clearly shows that a model that ignores the mRNA dynamics does not always capture the realistic distribution shape of the switching time distributions. In addition, we find that the holding time in the unbound state may have either a light-tailed or a heavy-tail distribution. According to both simulations and theory, we show that a heavy-tailed distribution tends to occur when the binding rate of protein to the gene or the transcription rate in the unbound gene state is very small compared to other rate constants.

Finally, we investigate the phase diagrams of distribution shapes for both the full and reduced models of an autoregulatory circuit. For negative feedback loops, the holding time in the unbound state generally has a bell-shaped distribution in the regime of slow switching, a bimodal distribution in the regime of relatively fast switching, and a decaying distributing in the regime of fast switching. Furthermore, we also find that as the transcription rate in the unbound state increases, a negative feedback loop undergoes a triphasic stochastic bifurcation from the decaying phase to the bimodal phase, and then to the bell-shaped phase. In contrast, for a positive feedback loop, the holding time in the unbound state has a decaying distribution in most cases; the system can produce a bell-shaped distribution only when the binding rate is very small and when the unbinding rate is controlled within a narrow range. For the reduced model, the bimodal parameter region disappears for negative feedback systems and the bell-shaped parameter region disappears for positive feedback systems.

For simplicity and analytical tractability, we here only derive the switching time distributions in an autoregulatory feedback loop. We anticipate that the results in the present paper can be generalized to more complex gene regulatory networks [26, 41]. In addition, while we have an implicit effective description of mRNA and protein dilution due to cell division, via the effective mRNA and protein decay rates, it has recently been shown that, in some parameter regimes, this type of model cannot capture the stochastic dynamics predicted by models with an explicit description of the cell cycle [42, 43]. We also hope to investigate the effects of cell growth and division on the switching time distributions.

## Appendices

### A. Analytical distribution of the unbound duration for the full model

In Eq. (4), we have shown that the generating function *H*(*x, y, t*) satisfies the first-order linear PDE

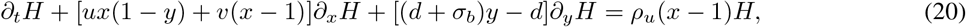

together with the initial condition

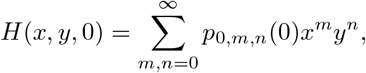

which can be determined by the initial distribution of the system. Recall that a first-order linear PDE can be solved using the method of characteristics. The characteristic curve (*x*(*t*), *y*(*t*)) associated with Eq. (20) is determined by following ODEs:

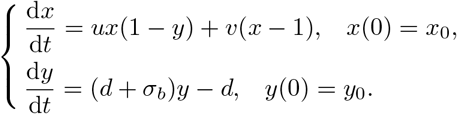

Solving these equations yields

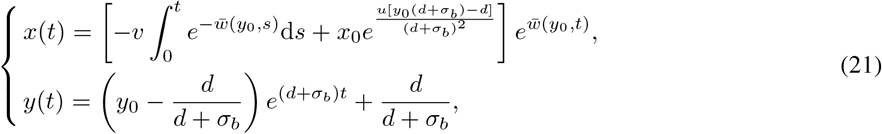

where

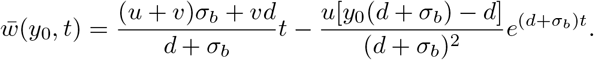

To proceed, we introduce the function 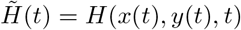, which is nothing but the value of the function *H* along the characteristic curve (*x*(*t*), *y*(*t*)). From Eq. (20), it is easy to see that 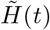 satisfies the following ODE:

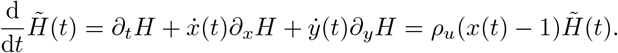

Solving this equation yields

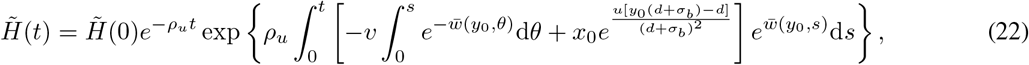

where 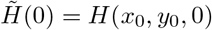 can be determined by the initial distribution of the system. To solve *H*(*x, y, t*), we need to determine the values of *x*_0_ and *y*_0_ such that the characteristic curve passes through (*x*_0_, *y*_0_) at time 0 and also passes through (*x, y*) at time *t*. From Eq. (21), it is easy to see that

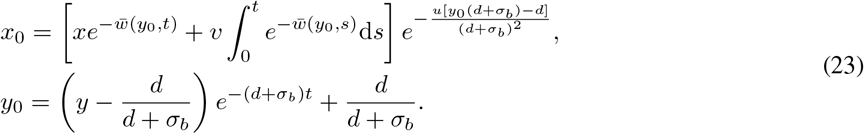

Finally, substituting Eq. (23) into Eq. (22) yields the analytical expression of *H*(*x, y, t*) given in Eq. (7).

### B. Derivation of the steady-state hitting distribution

In the main text, we have shown that when computing the unbound duration distribution in steady-state conditions, the initial condition should be chosen as the steady-state hitting distribution, i.e. the mRNA and protein number distribution just after switching from the bound to the unbound state after a sufficient number of switching events. To obtain the analytical expression of the hitting distribution, we introduce the following notation. Let *α*_*t*_ be the gene state in a single cell at time *t*, let *M*_*t*_ be the mRNA number at time *t*, and let *N*_*t*_ be the protein number at time *t*. Recall that the process

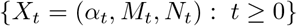

is a continuous-time Markov chain with state space *S* = {(*i, m, n*) : *i* = 0, 1, *m, n* ≥ 0}. The transition diagram of this Markov chain is illustrated in Fig. 1(b). Moreover, let *Q* = (*q*_*xy*_)_*x*,*y*∈*S*_ be the generator matrix of this Markov chain, where *x* and *y* denotes two microstates of the system. To proceed, let

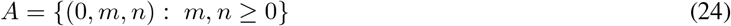

be the collection of all microstates in the unbound gene state and let

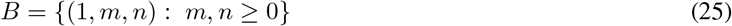

be the collection of all microstates in the bound gene state. We will next compute the steady-state hitting distribution of the set *A* by the process *X* [44].

To compute the steady-state hitting distribution, we first calculate the *k*th hitting distribution, i.e. the distribution when the system hits the set *A* for the *k*th time. Let

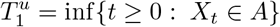

be the first hitting time of the set *A* and let

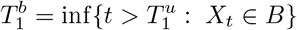

be the first hitting time of the set *B*. Then the first duration spent in the set *A* is given by 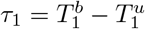. Similarly, let

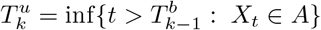

be the *k*th hitting time of the set *A* and let

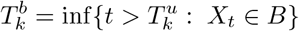

be the *k*th hitting time of the set *B*. Then the *k*th duration spent in the set *A* is given by 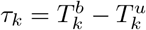. Then the *k*th hitting distribution of the set *A* by the process *X* is defined by

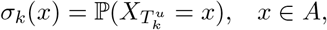

and the *k*th hitting distribution of the set *B* by the process *X* is defined by

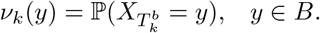

To compute the hitting distribution, we rewrite the generator matrix *Q* as the block form

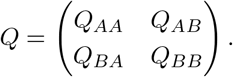

Using the classical theory of continuous-time Markov chains [44, 45], it is not difficult to prove that *ν*_*k*_ and *σ*_*k*_ are related by

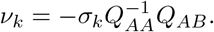

Similarly, it is easy to prove that *σ*_*k*+1_ and *ν*_*k*_ are related by

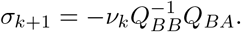

Combining the above two equations, we obtain

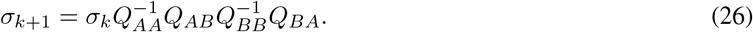

By the ergodic theorem, after a sufficient number of switching events, the *k*th hitting distribution *σ*_*k*_ will converge to the steady-state hitting distribution

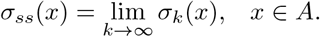

Taking *k* → ∞ in Eq. (26), we find that the steady-state hitting distribution *σ*_*ss*_(*x*) should satisfy

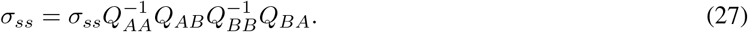

Let *π* = (*π*_*A*_, *π*_*B*_) = (*π*_*x*_)_*x*∈*S*_ denote the steady-state distribution of the process {*X*_*t*_ : *t* ≥ 0} and let **1** = (1, · · ·, 1)^*T*^ denote the column vector whose components are all 1. Since the steady-state distribution *π* satisfies *πQ* = 0, we obtain

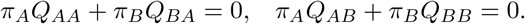

Finally, solving Eq. (27) yields

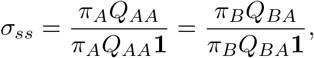

which can be written in components as

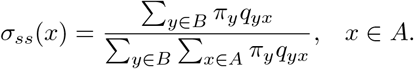

This indicates that the steady-state hitting probability of any microstate *x* ∈ *A* is exactly the total steady-state probability flux from the set *B* to this microstate, divided by the total steady-state probability flux from the set *B* to the set *A*.

### C. Steady-state joint distribution of mRNA and protein numbers for the full model

Here we derive the steady-state joint distribution of mRNA and protein numbers for the full model when the gene switches slowly between the two states, i.e. *σ*_*u*_, *σ*_*b*_ ≪ *ρ*_*u*_, *ρ*_*b*_, *u, v, d*. Note that when *σ*_*u*_ and *σ*_*b*_ are small, the microstates in the unbound gene state, i.e. the microstates in the set *A* defined in Eq. (24), and the microstates in the bound gene state, i.e. the microstates in the set *B* defined in Eq. (25) are both in rapid pre-equilibrium and hence can be aggregated into two groups (unbound and bound groups). Let 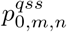 and 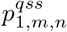 denote the quasi-steady-state distributions of mRNA and protein numbers for the unbound and bound groups, respectively. From Fig. 1(a), the evolution of the quasi-steady-state distribution 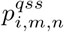 is given by

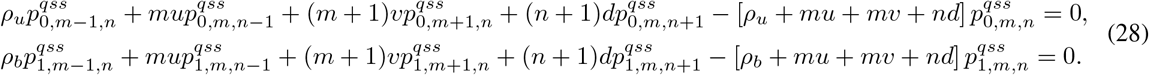

To proceed, let

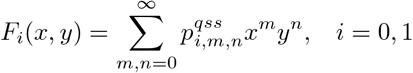

denote the generating functions of the quasi-steady-state distributions. Then Eq. (28) can be converted into the following system of PDEs:

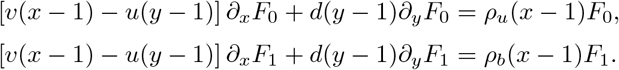

These PDEs have been solved exactly in [46, 47] and the analytical solution of *F*_*i*_(*x, y*) is given by Eq. (11) in the main text. Taking derivations of the generating functions *F*_*i*_(*x, y*) at *x* = *y* = 0 gives the quasi-steady-state distributions

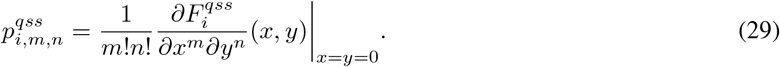

Since gene switching is very slow, according to the classical two-time-scale simplification techniques of Markov chains called averaging [48], the full model illustrated in Fig. 1(a) can be simplified to the following two-state system:

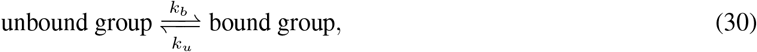

with effective transition rates *k*_*b*_ and *k*_*u*_. Since the transition from the bound microstate (1, *m, n*) to the unbound microstate (0, *m, n* + 1) occurs at constant rate *σ*_*u*_, the effective unbinding rate is also *σ*_*u*_, i.e. *k*_*u*_ = *σ*_*u*_. Moreover, since the transition from the unbound microstate (0, *m, n*) to the bound microstate (1, *m, n* − 1) occurs at rate *σ*_*b*_*n*, according to the averaging theory [48], the effective binding rate is given by

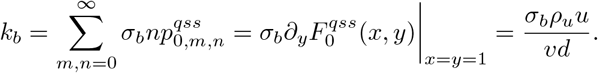

Hence the steady-state distribution for the two-state system given in Eq. (30) is given by

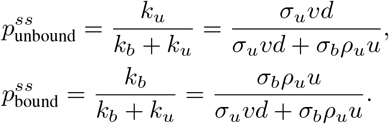

Finally, in the regime of slow switching, the steady-state joint distribution of mRNA and protein numbers is given by

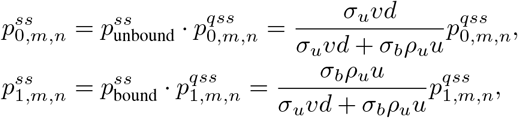

where 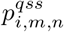 has been derived in Eq. (29).

## Acknowledgements

We thank Prof. Youming Li for valuable suggestions on the manuscript. C. J. acknowledges support from National Natural Science Foundation of China with NSAF grant No. U2230402 and grant No. 12271020.

