## Supplementary Material for "Exact switching time distributions for autoregulated gene expression models with mRNA and protein descriptions"

#### Contents

**1 Analytical distribution of the unbound duration for the reduced model** **2**

### 1 Analytical distribution of the unbound duration for the reduced model

We next compute the analytical unbound duration distribution for the reduced model shown in Fig. 1(c). Recall that the reduced model can be viewed as the limit of the full model illustrated in Fig. 1(a) as $\gamma = v/d \to \infty$ while keeping $u/v$ as constant. To proceed, let $\lambda = u + v$, $p = u/(u+v)$, and $q = 1 - p = v/(u+v)$. Note that $\gamma \to \infty$ while keeping $u/v$ as constant is equivalent to $\lambda \to \infty$ while keeping $p$ and $q$ as constant. Recall that for the full model, the generating function $H(x, y, t)$ has the following explicit expression:

$$\begin{aligned} H(x,y,t) &= H(\alpha(x,y,t), \beta(y,t), 0) \\ &\quad \times \exp\left[\rho_u \left(x \int_0^t e^{w(y,t,s)-w(y,t,t)} \mathrm{d}s + v \int_0^t \mathrm{d}s \int_s^t e^{w(y,t,s)-w(y,t,\theta)} \mathrm{d}\theta - t\right)\right]. \end{aligned} \tag{1}$$

We first consider the limit of $\lambda \to \infty$ in the domain $E = \{y \in \mathbb{R} : -1 < y \leq d/(d+\sigma_b)\}$. Note that

$$w(y,t,s) - w(y,t,t) = \frac{\lambda(\sigma_b + qd)}{d+\sigma_b}(s-t) - \frac{\lambda p[(d+\sigma_b)y - d]}{(d+\sigma_b)^2}\left(e^{-(d+\sigma_b)(t-s)} - 1\right).$$

It is clear that

$$\lim_{\lambda \to \infty} [w(y,t,s) - w(y,t,t)] = -\infty,$$

for any $t > s$ and $(d+\sigma_b)y \leq d$. Hence we have

$$\lim_{\lambda \to \infty} x \int_0^t e^{w(y,t,s)-w(y,t,t)} \mathrm{d}s = 0, \tag{2}$$

whenever $(d+\sigma_b)y \leq d$. On the other hand, note that

$$w(y,t,s) - w(y,t,\theta) = -\frac{\lambda(\sigma_b + qd)}{d+\sigma_b}(\theta - s) - \frac{\lambda p[(d+\sigma_b)y - d]}{(d+\sigma_b)^2} e^{-(d+\sigma_b)(t-\theta)}\left(e^{-(d+\sigma_b)(\theta-s)} - 1\right). \tag{3}$$

To proceed, let

$$\eta = \frac{\sigma_b + qd}{d+\sigma_b} - \frac{p[(d+\sigma_b)y - d]}{d+\sigma_b} e^{-(d+\sigma_b)(t-s)},$$

and let $\epsilon > 0$ be small enough. Obviously, we have $\eta > 0$ whenever $(d+\sigma_b)y \leq d$. Since $e^s = 1 + s + o(s)$ when $s$ is small, it is easy to see that

$$\begin{aligned} &\int_s^{s+\epsilon} \exp\left\{-\frac{\lambda(\sigma_b + qd)}{d+\sigma_b}(\theta - s) - \frac{\lambda p[(d+\sigma_b)y - d]}{(d+\sigma_b)^2} e^{-(d+\sigma_b)(t-\theta)}\left(e^{-(d+\sigma_b)(\theta-s)} - 1\right)\right\} \mathrm{d}\theta \\ &= \int_0^{\epsilon} \exp\left\{-\lambda\left[\frac{(\sigma_b + qd)}{d+\sigma_b}\theta + \frac{p[(d+\sigma_b)y - d]}{(d+\sigma_b)^2} e^{-(d+\sigma_b)(t-s)}\left(1 - e^{(d+\sigma_b)\theta}\right)\right]\right\} \mathrm{d}\theta \\ &\sim \int_0^{\epsilon} e^{-\lambda\eta\theta} \mathrm{d}\theta = \frac{1}{\lambda\eta}(1 - e^{-\lambda\eta\epsilon}). \end{aligned} \tag{4}$$

Moreover, we have

$$\begin{aligned} &\int_{s+\epsilon}^{t} \exp\left\{-\frac{\lambda(\sigma_b + qd)}{d+\sigma_b}(\theta - s) - \frac{\lambda p[(d+\sigma_b)y - d]}{(d+\sigma_b)^2} e^{-(d+\sigma_b)(t-\theta)}\left(e^{-(d+\sigma_b)(\theta-s)} - 1\right)\right\} \mathrm{d}\theta \\ &\leq \int_{\epsilon}^{t-s} e^{-\lambda\eta\theta} \mathrm{d}\theta = \frac{1}{\lambda\eta}(e^{-\lambda\eta\epsilon} - e^{-\lambda\eta(t-s)}), \end{aligned} \tag{5}$$

where we have used the fact that $e^s \geq 1 + s$. Combining Eqs. (3), (4), and (5), it is easy to see that

$$\lim_{\lambda \to \infty} \lambda q \int_s^t e^{w(y,t,s)-w(y,t,\theta)} \mathrm{d}\theta = \frac{q(d+\sigma_b)}{(\sigma_b + qd) - p[(d+\sigma_b)y - d]e^{-(d+\sigma_b)(t-s)}}, \tag{6}$$

whenever $(d+\sigma_b)y \le d$. It then follows from Eqs. (2) and (6) that

$$\begin{aligned}
&\lim_{\lambda\to\infty} \exp\left[\rho_u\left(x\int_0^t e^{w(y,t,s)-w(y,t,t)}\mathrm{d}s + \lambda q\int_0^t \mathrm{d}s\int_s^t e^{w(y,t,s)-w(y,t,\theta)}\mathrm{d}\theta - t\right)\right] \\
&= \exp\left[\rho_u\left(\int_0^t \frac{q(d+\sigma_b)}{(\sigma_b+qd)-p[(d+\sigma_b)y-d]e^{-(d+\sigma_b)(t-s)}}\mathrm{d}s - t\right)\right] \\
&= e^{-\rho_u t}\left\{\frac{(dq+\sigma_b)e^{(d+\sigma_b)t}-[(d+\sigma_b)y-d]p}{(d+\sigma_b)(1-py)}\right\}^{\frac{\rho_u q}{dq+\sigma_b}}.
\end{aligned} \tag{7}$$

Note that in the limit of $\lambda \to \infty$, mRNA molecules vanish most of the time. Hence we have $p_{0,m,n}(t) \approx \delta_{m,0}p_{0,n}(t)$, where $p_{i,m,n}(t)$ is the distribution of the full model and $p_{0,n}(t)$ is the distribution of the reduced model. This indicates that

$$\begin{aligned}
H(\alpha(x,y,t),\beta(y,t),0) &= \sum_{m,n=0}^{\infty} \delta_{m,0}p_{0,n}(0)\alpha(x,y,t)^m\beta(y,t)^n \\
&= \sum_{n=0}^{\infty} p_{0,n}(0)\beta(y,t)^n = H(\beta(y,t),0),
\end{aligned} \tag{8}$$

where the term $H(x,y,t)$ on the left-hand side represents the generating function for the full model and the term $H(y,t)$ on the right-hand side represents the generating function for the reduced model. Combining Eqs. (1), (7), and (8), for any $y \in E$, we have proved that the generating function $H(y,t)$ for the reduced model is given by

$$H(y,t) = H(\beta(y,t),0) \times e^{-\rho_u t}\left\{\frac{(dq+\sigma_b)e^{(d+\sigma_b)t}-[(d+\sigma_b)y-d]p}{(d+\sigma_b)(1-py)}\right\}^{\frac{\rho_u q}{dq+\sigma_b}}. \tag{9}$$

Since $\sum_{n=0}^{\infty} h(n,t) = \mathbb{P}(\tau_1 > t) < 1$, it is easy to see that the convergence radius of the power series given in Eq. (13) must be greater than or equal to 1. Hence $H(y,t)$ is holomorphic in the unit disk $D = \{y \in \mathbb{C} : |y| < 1\}$. Recall the uniqueness theorem of holomorphic functions claims that if two holomorphic functions $f(y)$ and $g(y)$ in the domain $D$ coincide on some set $E \subset D$ containing at least one limit point in $D$, then $f(y) \equiv g(y)$ for all $y \in D$. It is clear that the right-hand side of Eq. (9), as a function of $y$, is holomorphic in the domain $D$. Then the generating function $H(y,t)$ for the reduced model is given by Eq. (9) for all $y \in D$.
